# Faster model-based estimation of ancestry proportions

**DOI:** 10.1101/2024.07.08.602454

**Authors:** Cindy G. Santander, Alba Refoyo Martinez, Jonas Meisner

## Abstract

Ancestry estimation from genotype data in unrelated individuals has become an essential tool in population and medical genetics to understand demographic population histories and to model or correct for population structure. The `ADMIXTURE` software is a widely used model-based approach to account for population stratification, however, it struggles with convergence issues and does not scale to modern human datasets or the large number of variants in whole-genome sequencing data. Likelihood-free approaches optimize a least square objective and have gained popularity in recent years due to their scalability. However, this comes at the cost of accuracy in the ancestry estimates in more complex admixture scenarios. We present a new model-based approach, `fastmixture`, which adopts aspects from likelihood-free approaches for parameter initialization, followed by a mini-batch expectation-maximization procedure to model the standard likelihood. In a simulation study, we demonstrate that the model-based approaches of `fastmixture` and `ADMIXTURE` are significantly more accurate than recent and likelihood-free approaches. We further show that `fastmixture` runs approximately 30*×* faster than `ADMIXTURE` on both simulated and empirical data from the 1000 Genomes Project such that our model-based approach scales to much larger sample sizes than previously possible. Our software is freely available at https://github.com/Rosemeis/fastmixture.

## Introduction

For the past two decades, unsupervised ancestry estimation has been a crucial element in studies on human evolutionary genetics and genome-wide association studies [1, 2]. Estimating global ancestry proportions in unrelated individuals and corresponding ancestral allele frequencies has been a classic way of correcting for population structure and understanding demographic processes that have shaped the evolutionary history of modern populations. Ancestry estimation from genotype data emerged with the model-based clustering approach proposed with `STRUCTURE` [3], where individuals are proportionally assigned to an assumed number of ancestral populations. They modeled the probability of the observed genotype data given ancestry proportions and ancestral allele frequencies using a Bayesian approach. Due to scalability issues, the Bayesian approach was later replaced by maximum likelihood models, which were optimized using expectation-maximization (EM) and block relaxation algorithms, this includes the widely used software `ADMIXTURE` [4, 5].

In the era of big data, where ever-growing cohorts contain thousands of individuals with genotype data for millions of genetic variants, classic state-of-the-art model-based approaches, such as `ADMIXTURE` and `STRUCTURE`, fail to scale due to computational intractability. Over the past decade, there have been multiple attempts to scale unsupervised ancestry estimation. These efforts have primarily been rooted in likelihood-free approaches, with a few exceptions that have attempted to scale the standard model-based approach. Model-based and likelihood-free approaches have been shown to be connected within the framework of matrix factorization, where both approximate an observed genotype matrix by lower rank matrices under different assumptions and constraints [6]. In likelihood-free approaches, it is common to optimize a different least square objective using either non-negative matrix factorization (NMF) [7, 8] or alternating least square (ALS) [9, 10]. The SCOPE software [10] has gained increased popularity due to its efficient implementation of an ALS approach, which is well-suited for biobank-scale datasets. Meanwhile for the model-based approaches, a stochastic variational inference algorithm [11] and a neural network autoencoder [12] have also been proposed in recent times.

As large-scale cohorts begin to expand and include more cosmopolitan representations of individuals around the globe, obtaining accurate ancestry estimates has become a crucial task of modern genomics. This is essential to properly correct for population structure in genome-wide association studies and mitigate the bias in the over-representation of European-descent samples in current cohorts due to the ever-increasing focus on genomics for population health applications [13, 14]. While convergence within a reasonable time frame is highly desirable for these extensive datasets, recent attempts to improve scalability most often compromise accuracy in the ancestry estimation, as demonstrated here. The field still lacks a method that scales to larger datasets while being as accurate as the classic state-of-the-art approaches.

We introduce `fastmixture`, a novel model-based method for estimating ancestry proportions and ancestral allele frequencies in unrelated individuals. We leverage randomized singular value decomposition (SVD) for initializing the ancestry proportions and ancestral allele frequencies, followed by a mini-batch accelerated scheme to speed up the convergence of the EM algorithm. We demonstrate in an extensive simulation study and on real data that `fastmixture` significantly outperforms the original `ADMIXTURE` software in terms of speed while maintaining higher accuracy compared to recently developed approaches for ancestry estimation.

## Material and methods

We define a diallelic genotype matrix of N individuals and M variants or single-nucleotide polymorphisms (SNPs) as **G** *∈ {*0, 1, 2*}*^*N×M*^, which corresponds to the minor allele counts. We describe the estimation of ancestry proportions and ancestral allele frequencies with K ancestral sources as a low-rank matrix factorization problem such that **G** *≈* 2**QP**^*T*^, with **Q** *∈* [0, 1]^*N×K*^ and the constraint 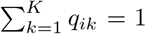, for i = 1, …, N, and **P** *∈* [0, 1]^*M×K*^. The matrix factorization problem can also be interpreted as the estimation of individual allele frequencies from the genotype matrix, **H** = **QP**^*T*^. Here, 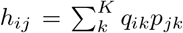 is the individual allele frequency of individual i at variant j can also be interpreted as the estimation of individual allele frequencies from the genotype matrix, assuming K ancestral sources. The individual allele frequency will correspond to the underlying parameter in a binomial sampling process of a genotype conditioned on population structure.

### Likelihood model

We estimate ancestry proportions and ancestral allele frequencies by maximizing the likelihood model introduced in `ADMIXTURE` [5]. The log-likelihood model is defined as follows, assuming independence for individuals and variants:

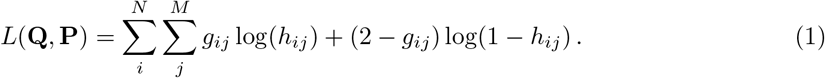

We utilize the expectation-maximization (EM) algorithm of `frappe` [4] and `ADMIXTURE` [5] to maximize the log-likelihood, where the EM updates at iteration t for entries of **Q** and **P** are defined as follows, respectively:

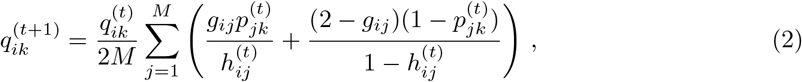

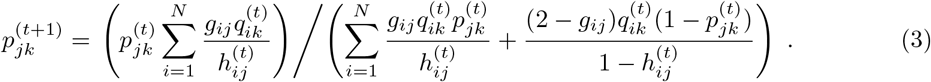

The EM algorithm is notorious for its slow convergence rate and to expedite this process, we employ a quasi-Newton (QN) acceleration scheme [15]. The QN acceleration scheme combines multiple EM updates into a larger jump in parameter space at the expense of an increased computational cost per iteration. More details on the acceleration scheme can be found in the supplementary material and Algorithm S1. The convergence criteria of the EM algorithm is defined by the difference in log-likelihood between two successive iterations:

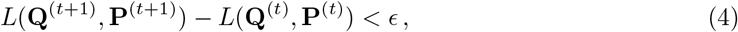

with ϵ being a user-defined threshold.

### SVD initialization

Multiple approaches for inferring population structure are connected under the umbrella of matrix factorization, each incorporating different conditions and constraints [6]. Therefore, we leverage the efficiency and speed of likelihood-free approaches to provide a better initialization of **Q** and **P** to aid the convergence rate of the EM algorithm in comparison to a standard random initialization.

We initialize **Q** and **P** using individual allele frequencies estimated from randomized singular value decomposition (SVD) performed on the genotype matrix, combined with an alternating least squares (ALS) approach. SVD is a widely used dimensionality reduction approach in population genetics, which infers continuous structure by extracting axes of genetic variation. The randomized SVD is defined as 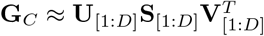s, which extracts the top D singular values and singular vectors, with **G**_*C*_ being the centered genotype matrix. The individual allele frequency of individual i at variant j is then approximated using the SVD as 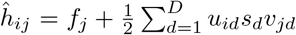, where f_*j*_ is the minor allele frequency of variant j. We use D = K *™* 1, as the top K *™* 1 singular vectors will capture the population structure of K distinct populations [16]. By initializing **P** randomly, we can then factorize the individual allele frequency matrix, 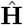, using an ALS approach to iteratively estimate both **Q** and **P**, which minimizes the following least square objective:

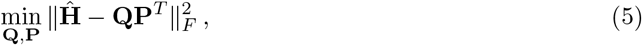

where ∥.∥_*F*_ is the Frobenius norm. The concept is similar to `ALStructure` [9] and `SCOPE` [10], which instead rely on latent subspace estimation for approximating the individual allele frequencies. The convergence criteria of the ALS approach is defined by the root mean square error (RMSE) of **Q** matrices between two successive iterations:

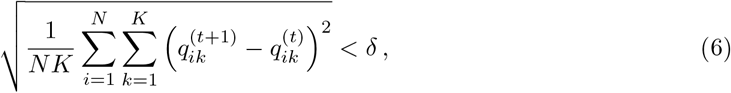

withδbeing a user-defined threshold. The initialization of **Q** and **P** is fully described in Algorithm 1.

#### Algorithm 1

`fastmixture` initialization

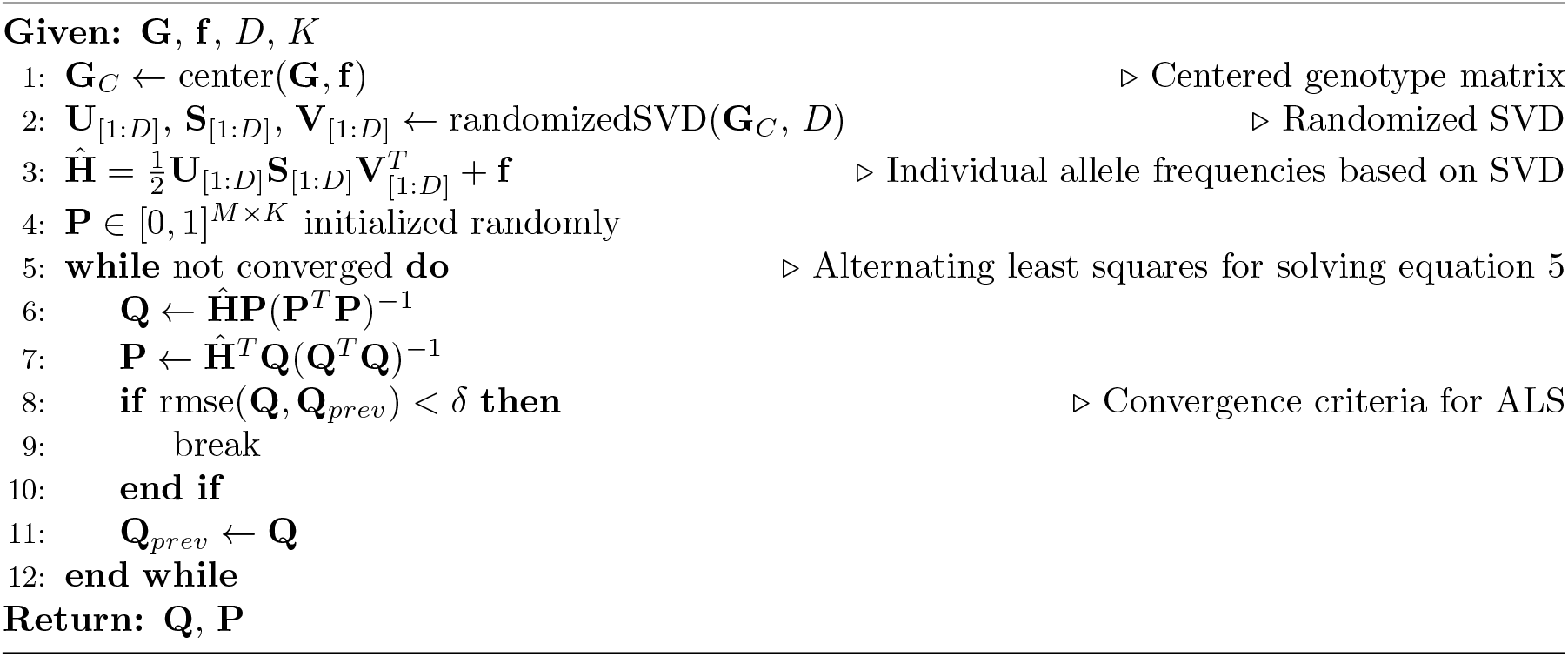

### Mini-batch optimization

To further accelerate the convergence rate of the EM algorithm, we introduce simple mini-batch updates inspired by stochastic gradient descent and stochastic EM algorithms [17]. In each iteration, we randomly split the M variants into B batches, and perform QN accelerated EM updates sequentially in each of the B batches. The strategy of sub-sampling variants was also explored in `TeraStructure` [11]. This results in the entries of **Q** being updated B times across the batches, while the entries of **P** are still only updated once per cycle. Following the mini-batch updates, full QN accelerated EM updates are applied to stabilize the parameters in every iteration. Given the stochastic nature of our mini-batch updates, we halve the number of batches B every time the log-likelihood estimate (Equation 1) fluctuates between iterations. The algorithm will therefore gradually converge towards a standard QN accelerated algorithm for B *→* 1. Our proposed minibatch setting resembles a mini-batch gradient descent approach, where the batch size is increased over time. For a detailed description of the `fastmixture` method, please refer to Algorithm 2.

#### Algorithm 2

`fastmixture` estimation

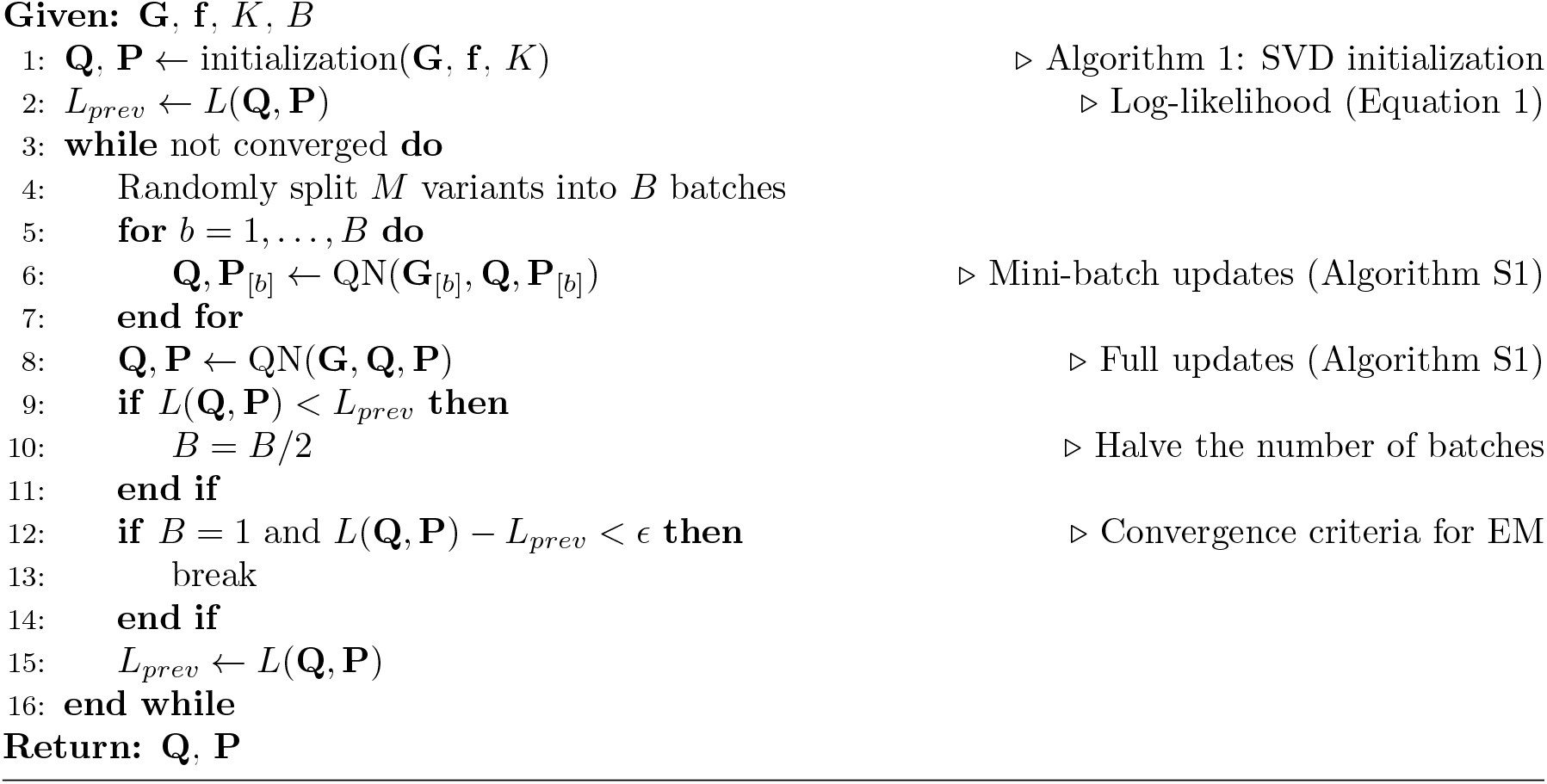

### Implementation details

The `fastmixture` software is implemented as a multithreaded command-line tool written in Python and Cython, which takes binary `PLINK` [18] files as input. It utilizes the NumPy library [19] for efficient array manipulation. We assume that the user has performed standard quality control and preprocessing, (e.g., variant filtering based on a minor allele frequency threshold and to only include unrelated samples). The genotype matrix is stored in an 8-bit integer format. For Algorithm 1, we read the centered genotypes in chunks to reduce the memory consumption and perform randomized SVD as introduced in PCAone [20]. This approach minimizes the memory footprint of `fastmixture`, which is primarily dominated by the genotype matrices of *MN* + 8*CN* bytes, with *C* being the chunk size of variants used in the randomized SVD. This allows our method to handle large-scale datasets effectively. The individual allele frequency matrix obtained in the randomized SVD, 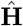, is computed implicitly through the singular matrices in the ALS approach to further reduce the memory footprint. Throughout the study, we use a starting point of *B* = 32 mini-batches to speed up the convergence of the EM algorithm, which works well across all the tested scenarios. We set the convergence criteria of the ALS approach and the EM algorithm to *δ* = 1.0 *×* 10^−4^ and ϵ = 0.5, respectively. Given the computational expense of the log-likelihood estimation step (Equation 1), which requires a full pass over the data, we only evaluate it every fifth iteration along with convergence checks to reduce the number of computations. If the entries of **Q** and **P** are out of domain during the ALS or accelerated EM updates, we simply map them back to their domain through truncation and projection procedures.

### Simulations

We simulate genotypes using the `msprime` [21] backwards-in-time coalescent model and infer true ancestral tracts using `tspop` [22]. To evaluate the capabilities of `fastmixture`, we assume different demographic models in four different scenarios, all featuring a single or multiple admixture events where the true ancestry proportions are known. In each scenario, we simulate a genetic segment of 250 Mb using a constant recombination rate of 1.28 *×* 10^−8^ [23] and a mutation rate of 2.36 *×* 10^−8^ [24]. We visualize the different demographic models in Figure 1A, Figure S3A, S1A, and S2A. A census event precedes the first admixture event in all simulations to track the true ancestry of the inherited segments in the sampled individuals. The lengths of the segments each individual has inherited from each source population are then aggregated to estimate the ground truth ancestry proportions using tspop. Scenarios A, B, and C are constructed from simple demographic models, where we assume a constant population size of 10,000 for all simulated populations, while Scenario D extends the out-of-Africa model [24] with an additional admixture event (American-Admixture) [25]. In Scenario A, B, and D, we sample 1,000 individuals, while in the more complex Scenario C, we sample 1,600 individuals. We perform standard filtering on minor allele frequencies at a threshold of 0.05, resulting in datasets consisting of 689,563 SNPs, 687,107 SNPs, 685,592 SNPs, and 500,114 SNPs for Scenario A, B, C, and D, respectively. An overview of the simulated datasets is provided in the supplementary material.

**Figure 1:**
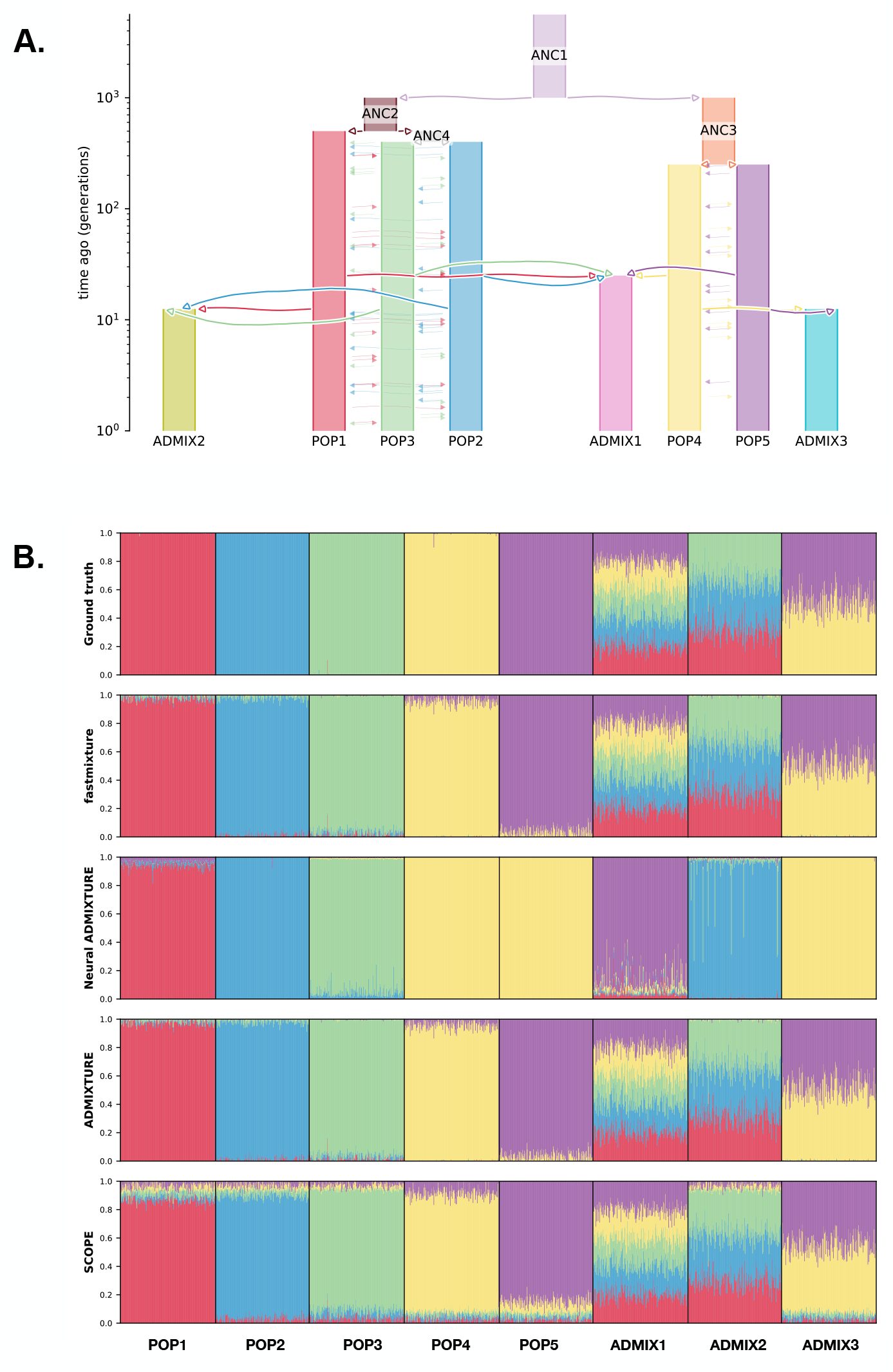
**A**. Demographic model of *Scenario C* with 200 individuals sampled from each of the eight populations. **B**. Admixture plots of the ancestry proportions in the simulated individuals for *K* = 5 with the ground truth at the top followed by the different software using their run with the highest log-likelihood.

### 1000 Genomes Project

We also evaluate our `fastmixture` software in the phase 3 release of the 1000 Genomes Project (1KGP) [26]. The dataset consists of genotype data of 2,504 individuals from 26 populations across the world, assigned to five super-populations: AFR (African ancestry), AMR (American ancestry), EAS (East Asian ancestry), EUR (European ancestry), SAS (South Asian ancestry). We keep diallelic SNPs with a minor allele frequency greater than the standard threshold of 0.05, resulting in a total of 6,864,700 SNPs. Due to computational complexity and runtime considerations for comparative analyses, we also construct a downsampled dataset, which we refer to as “1KGP Down”. This dataset is obtained by random downsampling or thinning the number of SNPs from the full dataset by a factor of ten, such that the downsampled dataset consists of 686,470 SNPs.

## Results

We evaluate and compare the performance of `fastmixture` (v0.93) against widely used software for estimating ancestry proportions: `ADMIXTURE` (v1.3.0) [5], Neural `ADMIXTURE` (v1.4.1) [12] and `SCOPE` [10]. All software are executed with their default parameters. `ADMIXTURE` and Neural `ADMIXTURE` are model-based approaches, like ours. `ADMIXTURE` maximizes the likelihood model using a block relaxation method combined with a similar quasi-Newton acceleration scheme to `fastmixture`, while `Neural ADMIXTURE` employs a neural network autoencoder approach. On the other hand, `SCOPE` is a likelihood-free approach that estimates the ancestry proportions and ancestral allele frequencies using an ALS method, differing from the maximum likelihood approaches of the other tools. In our simulation study, we assess these tools based on the root mean square error (RMSE) (Table 1) and the Jenson-Shannon divergence (JSD) (Table S2) between the estimated ancestry proportions and the ground truth ancestry proportions. We also report the log-likelihood estimates (Equation 1) (Table S3) for both the simulated and the empirical datasets of the 1000 Genomes Project. The different measures of assessment in the simulation study are further detailed in the supplementary material.

**Table 1:**
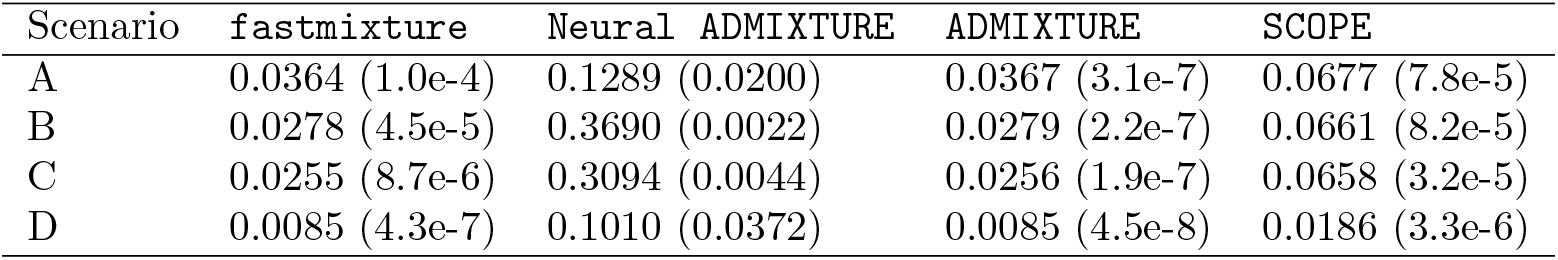
Root mean square error (RMSE) measures for estimated ancestry proportions in the four different simulation scenarios for the evaluated methods against the ground truth. The mean across five different runs is reported with the corresponding standard deviation in parenthesis.

Additionally, we compare the computational runtimes of `fastmixture` to the three other software (Figure 3 and Table S1). Notably, `ADMIXTURE` exhibits significant scalability issues, being approximately 30 times slower than `fastmixture` across all evaluated datasets. For instance, `ADMIXTURE` needs more than 40 hours to complete a single run for *K* = 5 on the full 1KGP dataset. In contrast, `fastmixture` has comparable runtimes to the two other faster approaches in the simulation study. `SCOPE` is the fastest of all evaluated approaches, showcasing the appealing choice of optimizing the least square objective, a step also used for parameter initialization in `fastmixture`.

### Ancestry estimation in simulation studies

Under the simple demographic model of Scenario A (Figure S1), `ADMIXTURE` and `fastmixture` were the two most accurate software followed by SCOPE. When only considering the scalable methods, `fastmixture` clearly outperformed `SCOPE` in terms of RMSE (Table 1) and JSD (Table S2), where `SCOPE` appeared to produce highly noisy ancestry proportions in individuals of the unadmixed populations. On all accounts, Neural `ADMIXTURE` severely underperformed in estimating the ancestry proportions of the admixed individuals. Notably when admixture was introduced from more than three sources, as in Scenario B (Figure S2), Neural `ADMIXTURE` failed to detect POP4 as a separate population from POP3 and modeled the ADMIX population incorrectly as a separate unadmixed population. For Scenario B, both `ADMIXTURE` and `fastmixture` were again more accurate than SCOPE, while `fastmixture` was *∼*34*×* faster than `ADMIXTURE` (Figure 3 and Table S1). The noise introduced by `SCOPE` only increased for *K* = 4 in comparison to *K* = 3 in the simpler Scenario A.

We further evaluated the different software in a more complex simulation scenario, Scenario C, which includes five ancestral sources (K = 5) with symmetric migration patterns and three admixed populations (Figure 1). Consistent with results from the simpler scenarios, `fastmixture` and `ADMIXTURE` outperformed the two other approaches, with `fastmixture` being *∼*28*×* faster than `ADMIXTURE`. Due to the increased complexity of the simulation scenario, `SCOPE` exhibited an even greater increase of noise in its ancestry estimates, while `Neural ADMIXTURE` again failed to detect one of the unadmixed population sources and modeled two admixed populations as ancestral sources. Examining the accuracy of the ancestry estimates in each of the eight populations, we observed that `fastmixture` and `ADMIXTURE` performed similarly across the unadmixed and admixed populations, whereas `SCOPE` inferred more accurate ancestry estimates in the admixed populations in comparison to the unadmixed populations (Table S4 and S5).

Using the American-Admixture demographic model in Scenario D (Figure S3), we still consistently observed that `ADMIXTURE` and `fastmixture` perform similarly in terms of accuracy and log-likelihood, outputting results closest to the ground truth (Table 1 and Table S2), with `fastmixture` being *∼*28*×* faster than `ADMIXTURE`. While across all scenarios `SCOPE` exhibited the fastest runtimes, it also consistently underperformed in accuracy as measured with RMSE and JSD, however, markedly better than Neural `ADMIXTURE`.

### Testing hyperparameters

The number of initial batches in `fastmixture`, used for its mini-batch optimization, is a hyper-parameter. We tested the effect of changing the number of mini-batches in the more complex simulation scenario, Scenario C, having multiple admixture events and five source populations. We utilized B = *{*8, 16, 32, 64, 128*}*, including the default choice of *B* = 32, and reported the computational runtimes, log-likelihoods, RMSE, and JSD measures. Our results showed that `fastmixture` was robust to changes in *B*, as all evaluated choices consistently captured the same solutions with highly comparable assessment measures (Figure S4 and Table S6). Based on these findings, we conclude that *B* = 32 was an optimal choice, balancing both fast runtimes and highly accurate ancestry estimations.

We further evaluated the effectiveness of our SVD initialization by comparing it to random parameter initialization inside the `fastmixture` framework for Scenario C. We reported computational runtimes, log-likelihoods, RMSE, and JSD measures in Table S7, where the two initializations performed similarly but the SVD initialization approximately halves the runtime on average in comparison to having a random initialization. Therefore, our observed runtime gains relative to `ADMIXTURE` could largely be attributed to our proposed mini-batch optimization.

### Robustness to model misspecification

For most scenarios in ancestry estimation, the true number of ancestral sources is rarely known. We, therefore, tested and compared all software and their capabilities to deal with model misspecifications related to the number of ancestral sources used for Scenario C, which had a ground truth of *K* = 5. Here we would expect the ancestry estimations to capture older events in the demographic model for *K* < 5. The results comparing the software for *K* = *{*2, 3, 4*}* are displayed in Figures S6, S7 and S8, respectively, and their corresponding log-likelihoods are reported in Table S8. We note that `ADMIXTURE` only found the optimal solution in four out of five runs, thus showcasing its vulnerabilities due to random parameter initialization and a standard optimization approach. Due to the increased complexity of the simulation scenario, `SCOPE` exhibited an even further increased noise level in its ancestry estimates across all three values of K.

### Ancestry estimation in the 1000 Genomes Project

In addition to our simulation study, we also applied `fastmixture` to empirical data of the 1000 Genomes Project (1KGP), using both a downsampled version and the full set of variants. Specifically, the downsampled version was the only way to properly assess the performance of `ADMIXTURE` over multiple runs due to scalability issues (Figure S9). Here we observed, as in the simulation study, that `fastmixture` and `ADMIXTURE` performed comparably and the two methods achieved the highest log-likelihoods, followed by `SCOPE` and then Neural `ADMIXTURE` (Table S3), even though Neural `ADMIXTURE` maximizes the log-likelihood in its optimization approach.

For the full 1KGP dataset, we only performed a single run for `ADMIXTURE` due to its excessive computational runtime of > 40 hours, in comparison to the other software with runtimes of < 2 hours (Figure 3 and Table S1). Here `fastmixture` was *∼*30*×* faster on average than the `ADMIXTURE` run. We demonstrated again that `fastmixture` was the best performing method in terms of achieving the highest log-likelihood (Figure 2). Strikingly, Neural `ADMIXTURE` failed to accurately distinguish ancestral contributions in the AMR and SAS super-populations in both the full dataset and in 1KGP Down. Moreover, we saw a correlation of *r*^2^ = 0.72 between the estimated ancestry proportions in the downsampled and the full dataset for Neural `ADMIXTURE`, indicating poor robustness. This is in strong contrast to `fastmixture`, `ADMIXTURE` and `SCOPE` where the results between the full dataset and the downsampled one were highly correlated (*r*^2^ *≈* 1). We observed the same pattern of increased noise in the estimated ancestry proportions of the unadmixed individuals from SCOPE, which was consistent across the two 1KGP datasets.

**Figure 2:**
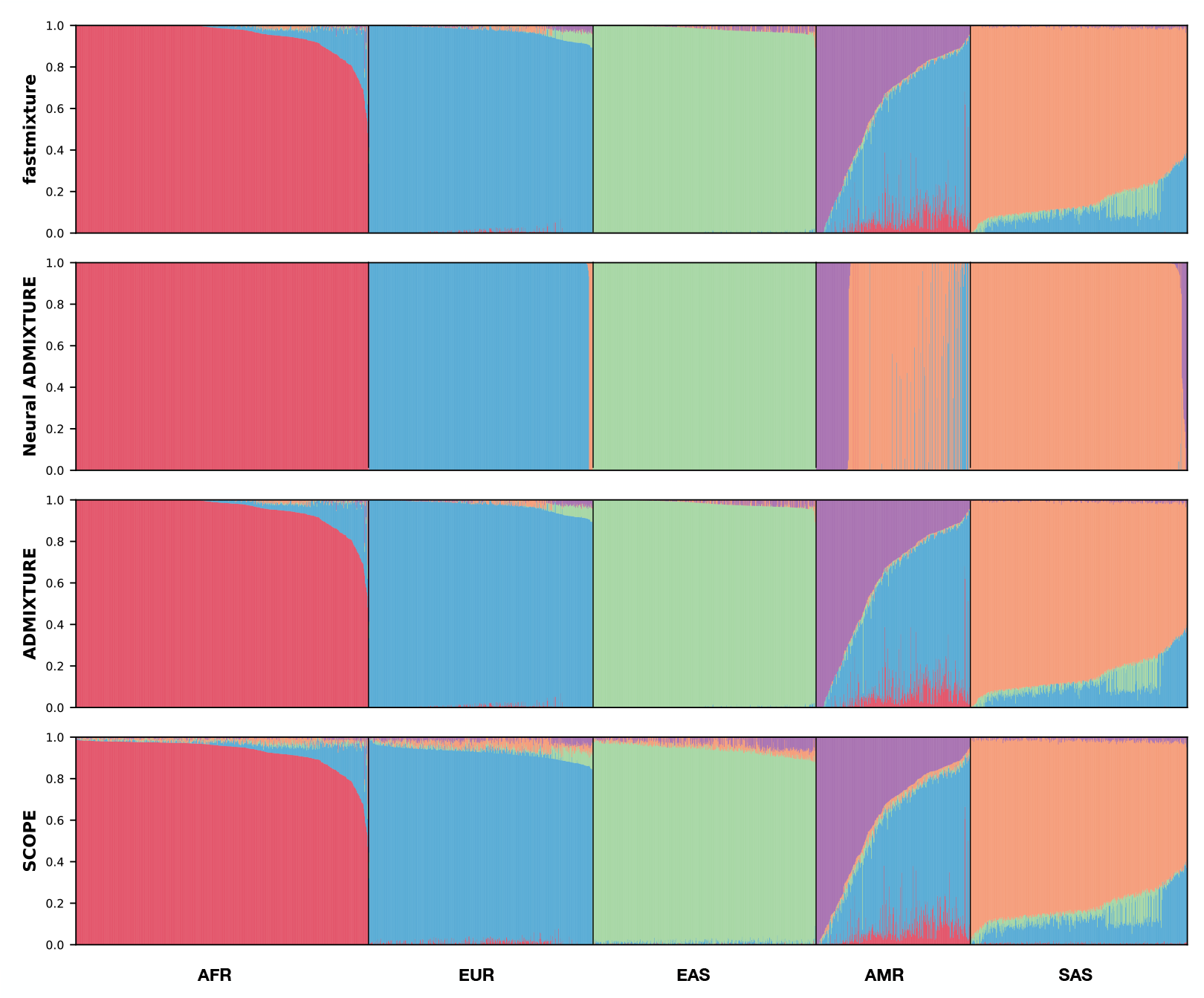
Admixture plots of the estimated ancestry proportions for *K* = 5 in the full version of the 1000 Genomes Project dataset (1KGP) by the different software using their run with the highest log-likelihood. AFR: African, EUR: European, EAS: East Asian, AMR: American, and SAS: South Asian ancestry.

**Figure 3:**
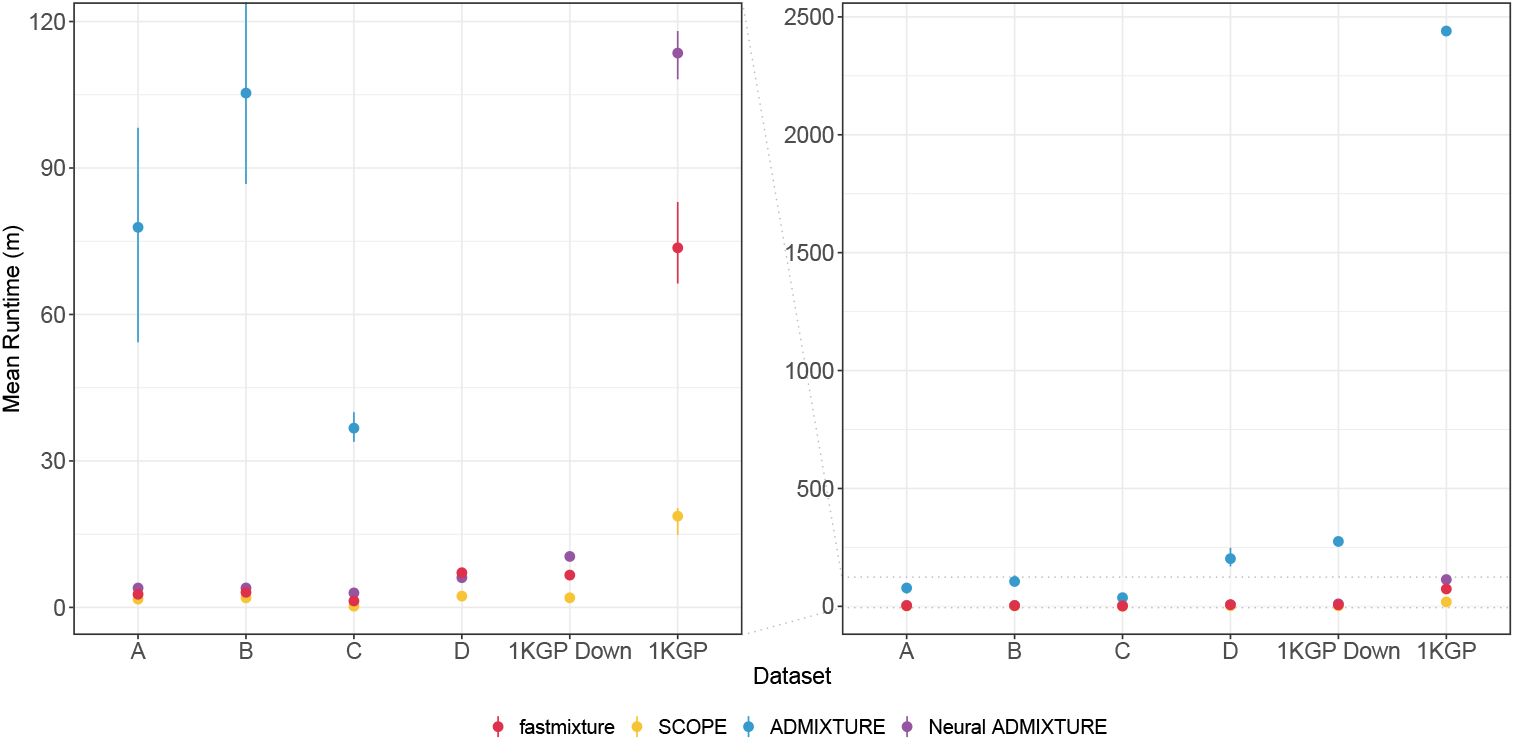
Runtime (in minutes) for the different software measured across five runs in the four simulated scenarios (A, B, C, and D), and in the two 1000 Genomes Project datasets: one including all individuals (1KGP), and the downsampled dataset (1KGP Down). The intervals display the full range of runtimes across the six runs. On the left, a zoomed-in version highlights the runtime differences between the software with more similar times. The results are also described in Table S1. All analyses have been performed on a cluster with an Intel(R) Xeon(R) Gold 6152 CPU using 64 threads.

## Discussion

We have presented, `fastmixture`, our novel method and software for ancestry estimation in genotype data using our extended model-based approach. We demonstrate that our approach performs comparably to `ADMIXTURE` while being *∼*30*×* faster on average across all evaluated datasets. Among the four methods assessed, `ADMIXTURE` and `fastmixture` stand out as the most accurate approaches as shown in our simulation study. In general, `fastmixture` estimates fast and accurate solutions that are robust to changes in model parameters and hyperparameters, such as initialization and number of initial mini-batches, which can be attributed to both its SVD initialization and accelerated mini-batch optimization procedure. It is well known that `ADMIXTURE` has scalability issues for large sample sizes and whole-genome sequencing data, which we also demonstrate using the 1KGP datasets. The results of `ADMIXTURE` are only based on a single seed in the full 1KGP SNP set due to a prohibitively excessive computational runtime of more than 40 hours, in contrast to *∼*74 minutes using `fastmixture`. We have therefore evaluated all software on a downsampled version as well, with 10*×* less SNPs, which is a common procedure in population genetic studies.

fastmixture has runtimes comparable to the recently introduced Neural `ADMIXTURE` software, which employs an autoencoder framework for speeding up the ancestry estimation. Note that the testing of the software has exclusively been conducted in a CPU-based setup. However, Neural `ADMIXTURE` struggles across all evaluated datasets and consistently performs the poorest among the evaluated methods, as it can only manage to model unadmixed populations in simpler demographic models. In Scenario *B* and C, Neural `ADMIXTURE` fails to detect unadmixed populations and models admixed populations incorrectly as ancestral sources. `SCOPE` emerges as the fastest approach across all evaluated datasets, demonstrating excellent scalability. However, its optimization of a simpler least squares objective compromises its ability to accurately estimate ancestry proportions, where it can be difficult to distinguish real admixture signals from noise.

When optimizing a different objective, it is expected that the log-likelihood estimates for `SCOPE` would be lower compared to model-based approaches. However, based on the RMSE and JSD measures against the ground truth in the simulation study, our results showcase that the least square objective used in likelihood-free approaches, such as SCOPE, is not an optimal choice in comparison to the likelihood model. Our findings suggest that the added noise in the ancestry proportions estimated in `SCOPE` are likely to increase further in scenarios with a larger *K* or more complex demographic models, as demonstrated in Scenario C. This limits the utility of `SCOPE` in association studies and precision medicine. Furthermore, we observed a general trend of major optimization issues for the Neural `ADMIXTURE` software across all evaluated scenarios. A critical issue appears to be in their convergence evaluation, where log-likelihood estimates are normalized across individuals and variants in their mini-batch training setup, causing premature convergence. This premature convergence negatively impacts their performance, despite Neural `ADMIXTURE` erroneously reporting faster runtimes than it would achieve at optimal solutions.

While our `fastmixture` software does not entirely solve the scalability issues of model-based approaches, it represents a significant step by enabling researchers to estimate accurate ancestry proportions for much larger sample sizes and whole-genome sequencing data. It will also facilitate a more feasible exploration of increased numbers of ancestral sources. We anticipate that `fastmixture` will be the preferred alternative to `ADMIXTURE` in future population genetic studies, and it will also allow researchers to correct for population structure in genome-wide association studies of moderate sample sizes, leveraging our accurate estimates of ancestry proportions.

## Data availability

The study uses genotype data from the phase 3 release of the 1000 Genomes Project [26], publicly available at http://ftp.1000genomes.ebi.ac.uk/vol1/ftp/release/20130502/. The results (trees and ancestry estimations) and processed genotype data files used in the study are available on Zenodo (https://doi.org/10.5281/zenodo.12683371) for reproducibility.

## Code availability

The `fastmixture` software is open-source and freely available on GitHub (https://github.com/Rosemeis/fastmixture). Scripts for reproducing the entire simulation study are available on Zenodo (https://doi.org/10.5281/zenodo.12683371).

## Author contributions

J.M. has conceived the study and implemented the method. C.G.S., A.R.M., and J.M. have performed the analyses, discussed the results, and contributed to the final manuscript.

## Acknowledgements

J.M. is supported by a grant from the Lundbeck Foundation (R380-2021-1225).

## Supplementary material

### Quasi-Newton acceleration scheme

We here describe the quasi-Newton scheme [15] used to accelerate the expectation-maximization (EM) algorithm of `fastmixture`. For simplicity and speed, we only utilize one secant condition (q = 1) in the scheme for both the mini-batch and the full EM updates such that the update rule is a combination of two standard EM steps. We define **Q**^(1)^ and **P**^(1)^ as the updated matrices after one standard EM step, using Equation 2 and Equation 3, respectively, and **Q**^(2)^ and **P**^(2)^ after two successive standard EM steps. The accelerated update rules are then described as follows and we refer to the original paper on the quasi-Newton scheme for more details:

#### Algorithm S1

quasi-Newton acceleration

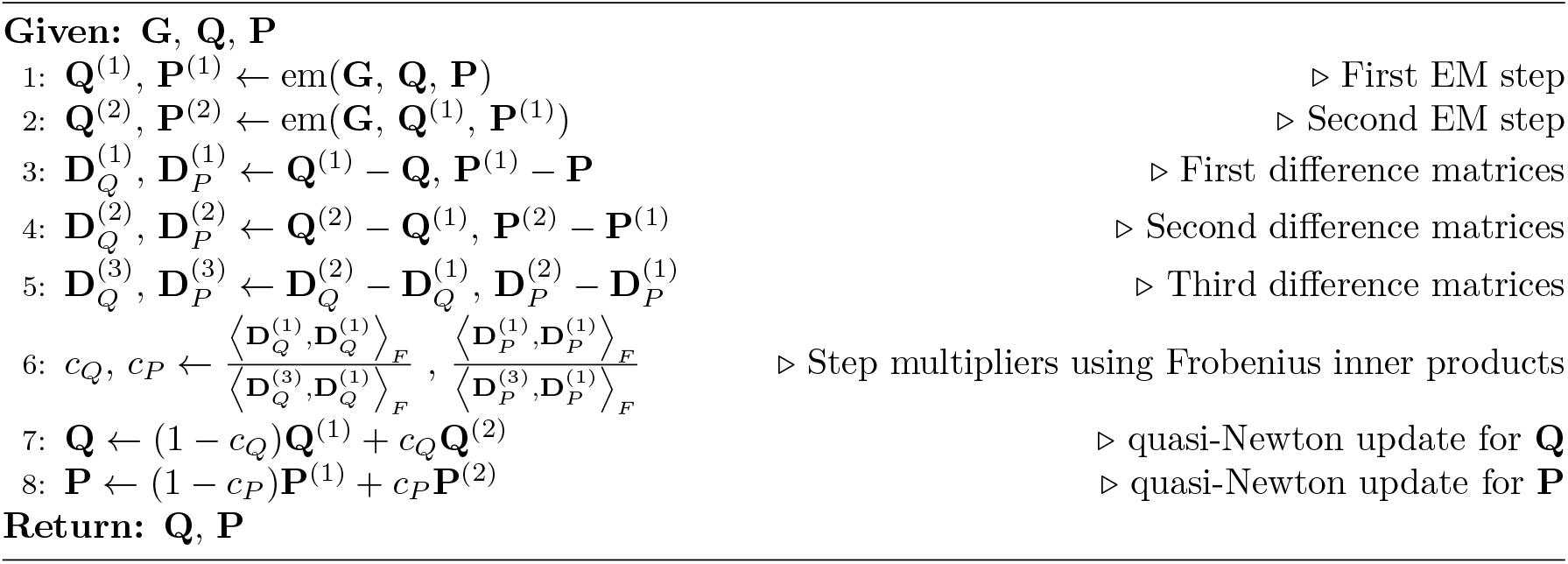

### Assessment measures

We here describe the different assessment measures used in the study. The log-likelihood estimate is already defined in Equation 1. **Q** refers to the estimated ancestry proportions and **Q**^*\**^ refers to the simulated ground truth ancestry proportions.

### Root mean square error (RMSE)

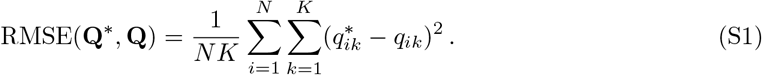

### Jensen-Shannon divergence (JSD)

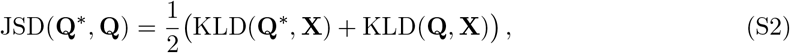

where 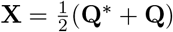 and KLD(*·, ·*) is the Kullback–Leibler divergence:

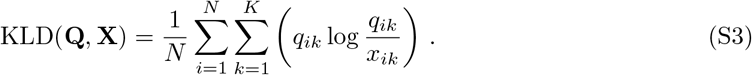

### Simulated data overview

All demographic models and scripts for reproducing the simulation scenarios are available on Zenodo (https://doi.org/10.5281/zenodo.12683371).

#### Common msprime options

- Segment length: 250 Mb
- Mutation rate: 2.36 *×* 10^−8^
- Recombination rate: 1.28 *×* 10^−8^

#### Scenario A

– *N* = 1,000 (250 individuals from each of the 4 simulated populations)
– *M* = 689,563
– 3 ancestral populations (POP1, POP2, POP3)
– 1 admixed population 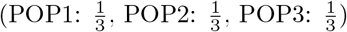

#### Scenario B

– *N* = 1,000 (200 individuals from each of the 5 simulated populations)
– *M* = 687,107
– 4 ancestral populations (POP1, POP2, POP3, POP4)
– 1 admixed population 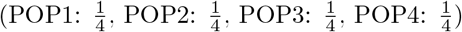

#### Scenario C

– *N* = 1,600 (200 individuals from each of the 8 simulated populations)
– *M* = 685,592
– 5 ancestral populations (POP1, POP2, POP3, POP4, POP5)
– 1 admixed population 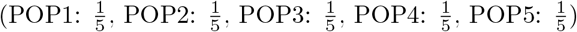
– admixed population 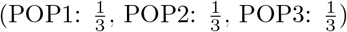
– admixed population 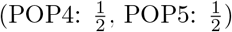

#### Scenario D

– *N* = 1,000 (250 individuals from each of the 4 simulated populations)
– *M* = 500,114
– 3 ancestral populations (AFR, EUR, EAS)
– 1 admixed population 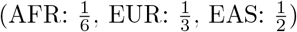

## Supplementary figures and tables

**Figure S1:**
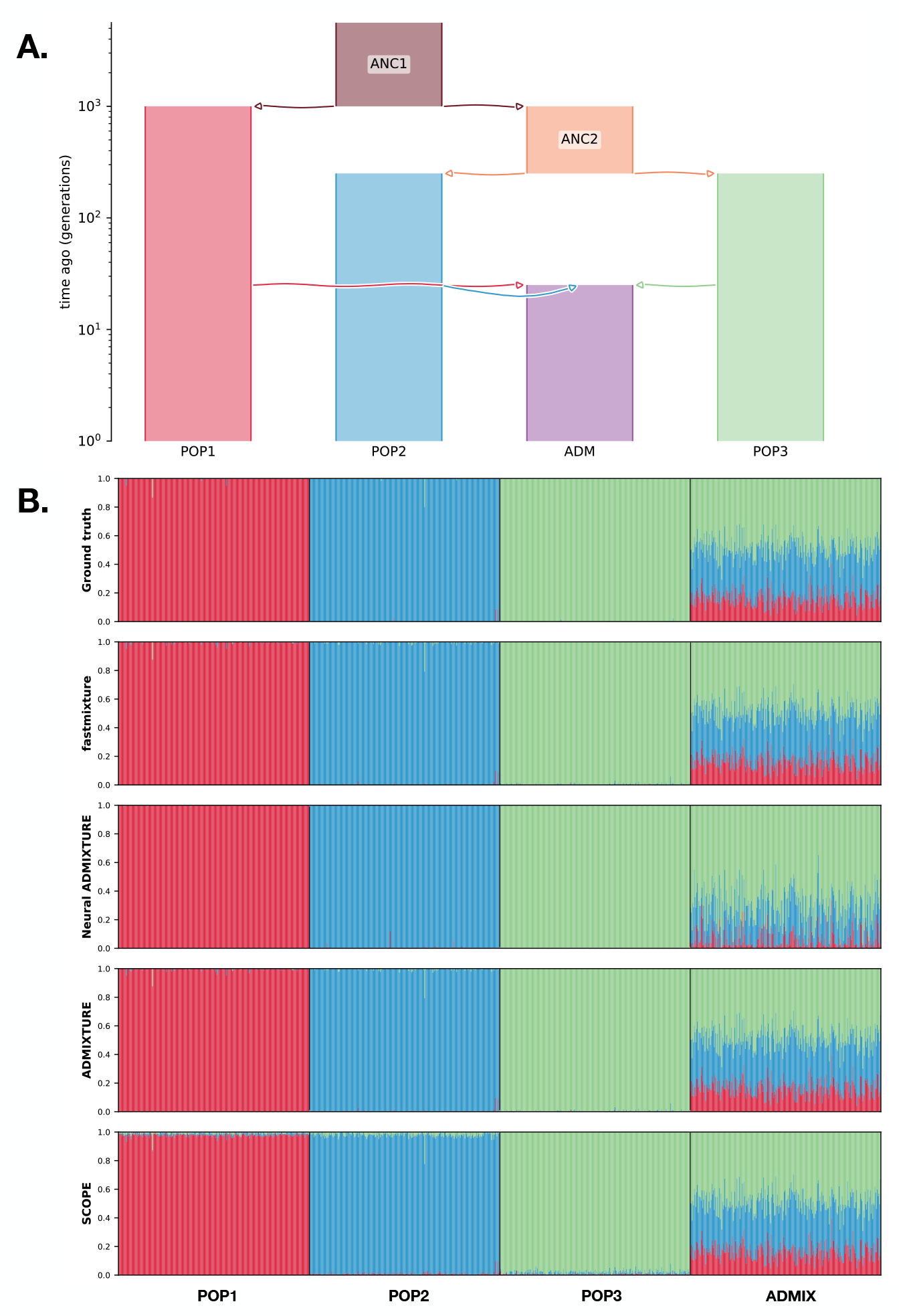
**A**. Demographic model of *Scenario A* with 250 individuals sampled from each of the four populations. **B**. Admixture plots of the ancestry proportions in the simulated individuals for *K* = 3 with the ground truth at the top followed by the different software using their run with the highest log-likelihood.

**Figure S2:**
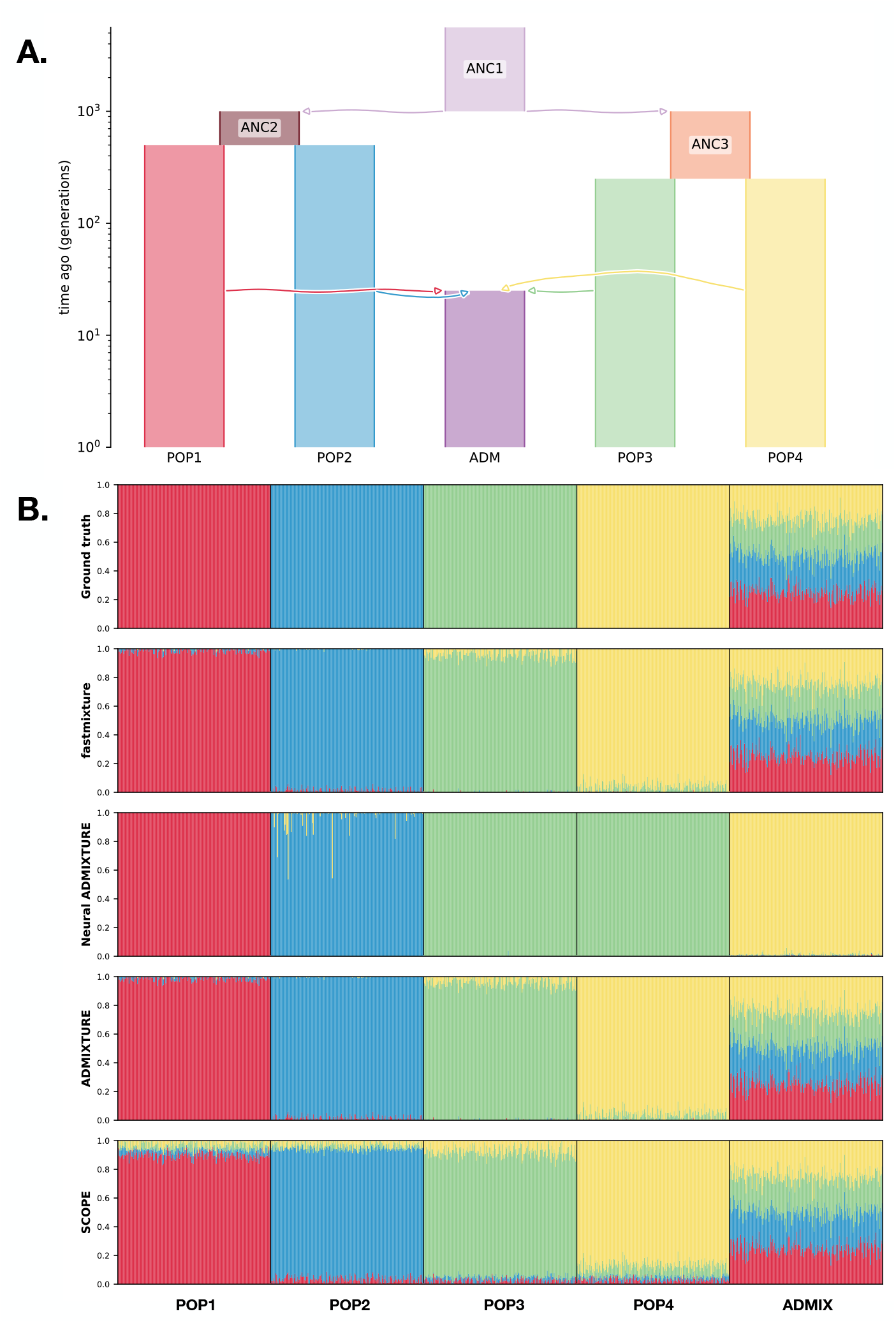
**A**. Demographic model of *Scenario B* with 200 individuals sampled from each of the five populations. **B**. Admixture plots of the ancestry proportions in the simulated individuals for *K* = 4 with the ground truth at the top followed by the different software using their run with the highest log-likelihood.

**Figure S3:**
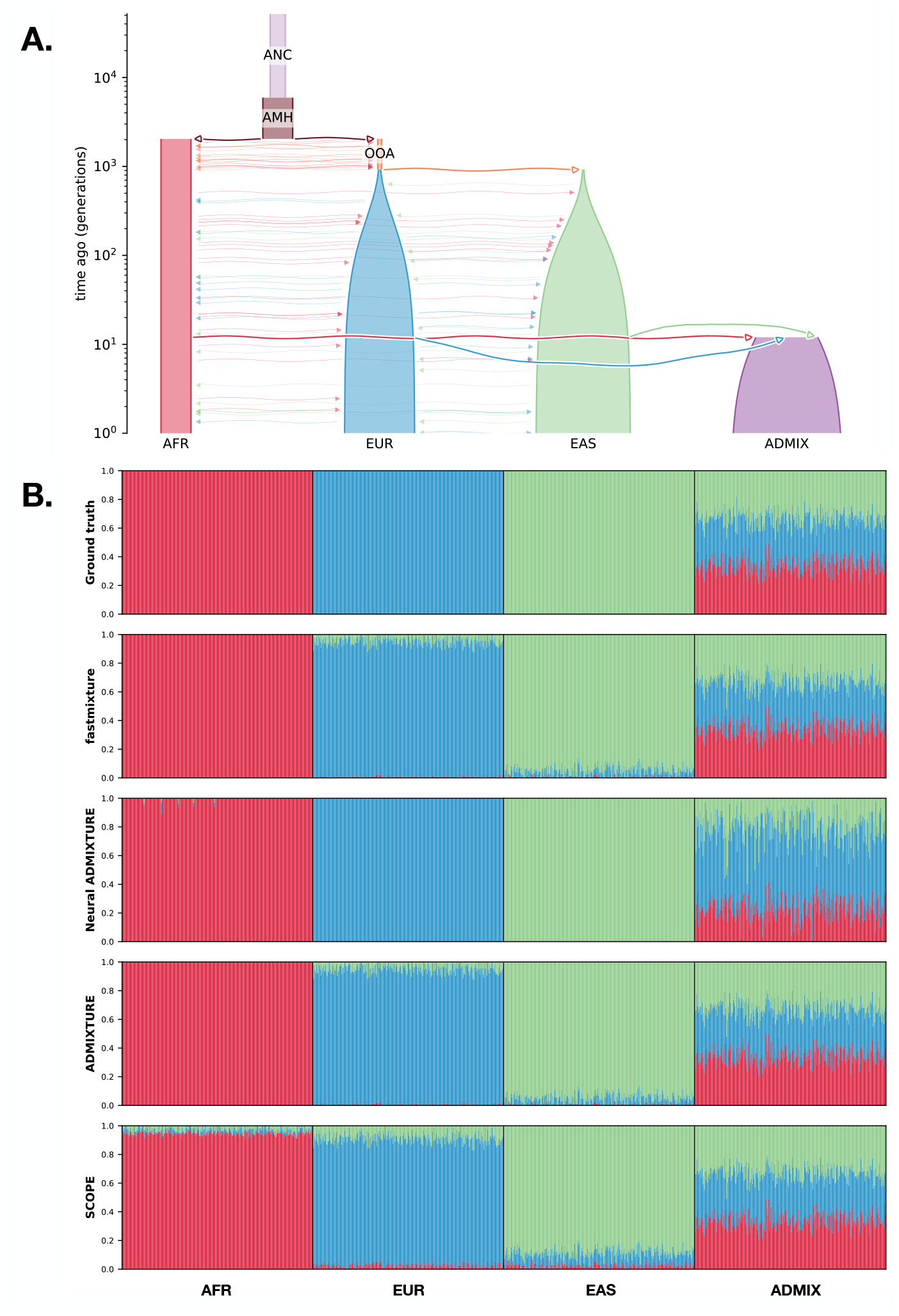
**A**. Demographic model of *Scenario D* (American admixture [25]) with 250 individuals sampled from each of the four populations. **B**. Admixture plots of the ancestry proportions in the simulated individuals for *K* = 3 with the ground truth at the top followed by the different software using their run with the highest log-likelihood.

**Figure S4:**
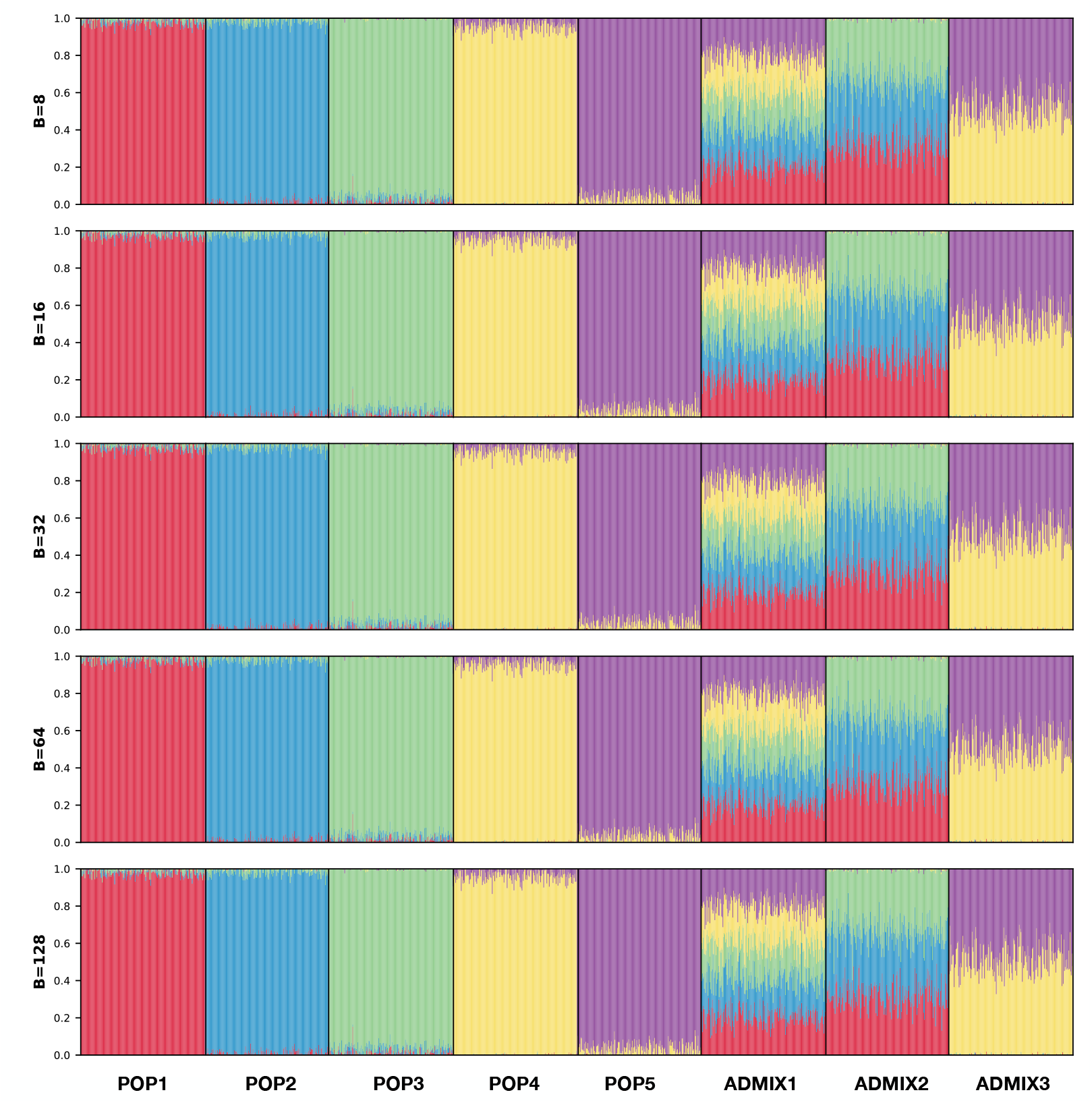
Admixture plots of the ancestry proportions in the simulated individuals for *K* = 5 of *Scenario C* by `fastmixture` with different batch sizes for *B* = *{*8, 16, 32, 64, 128*}* using their run with the highest log-likelihood.

**Figure S5:**
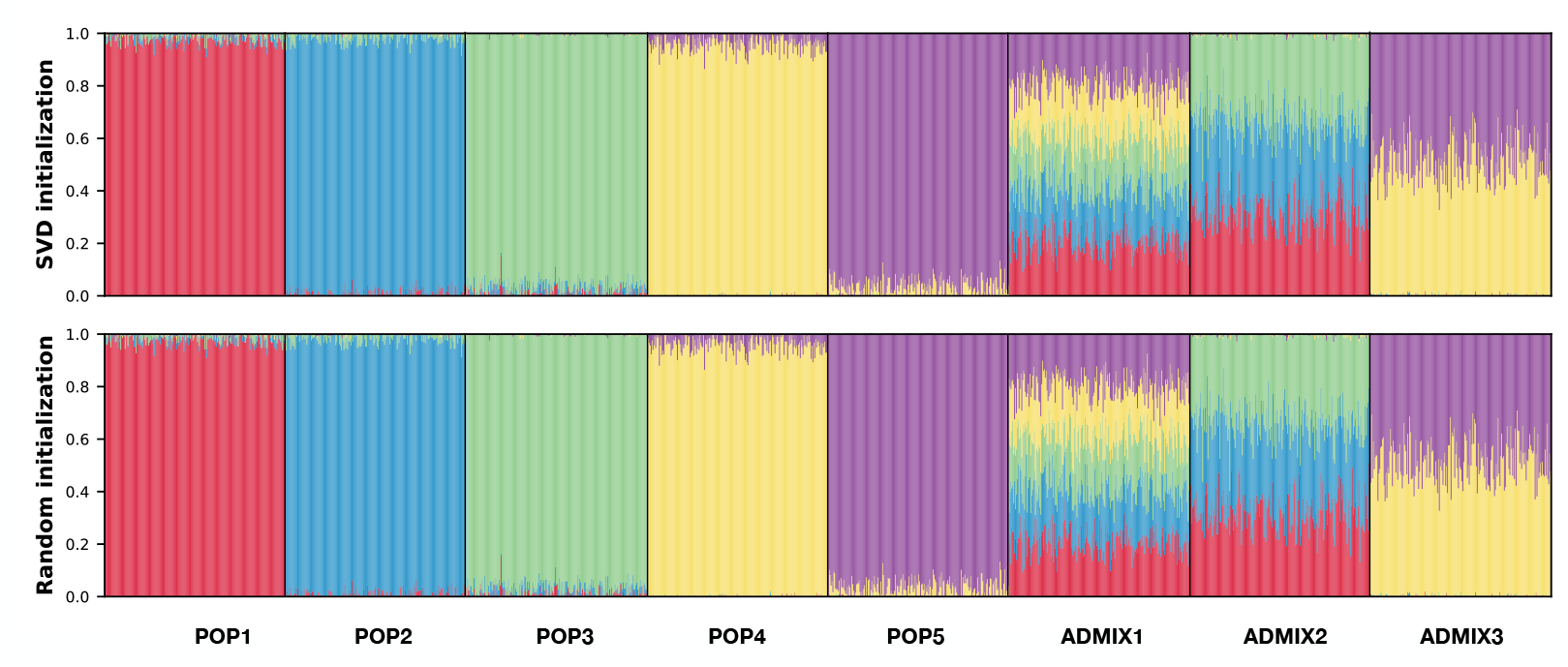
Admixture plots of the ancestry proportions in the simulated individuals for *K* = 5 of *Scenario C* by `fastmixture` with different types of initialization using their run with the highest log-likelihood.

**Figure S6:**
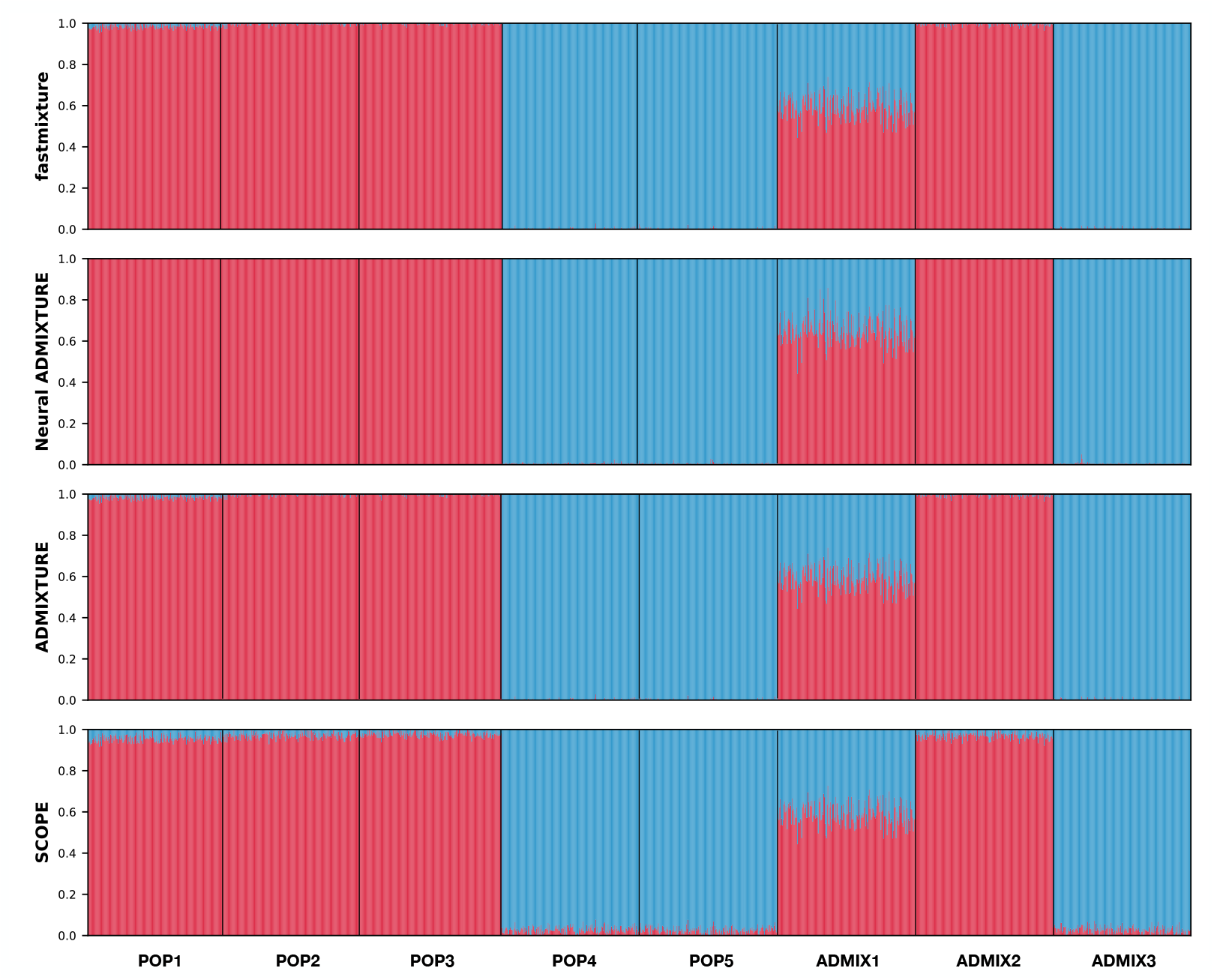
Admixture plots of the ancestry proportions in the simulated individuals for *K* = 2 of *Scenario C* by the different software using their run with the highest log-likelihood.

**Figure S7:**
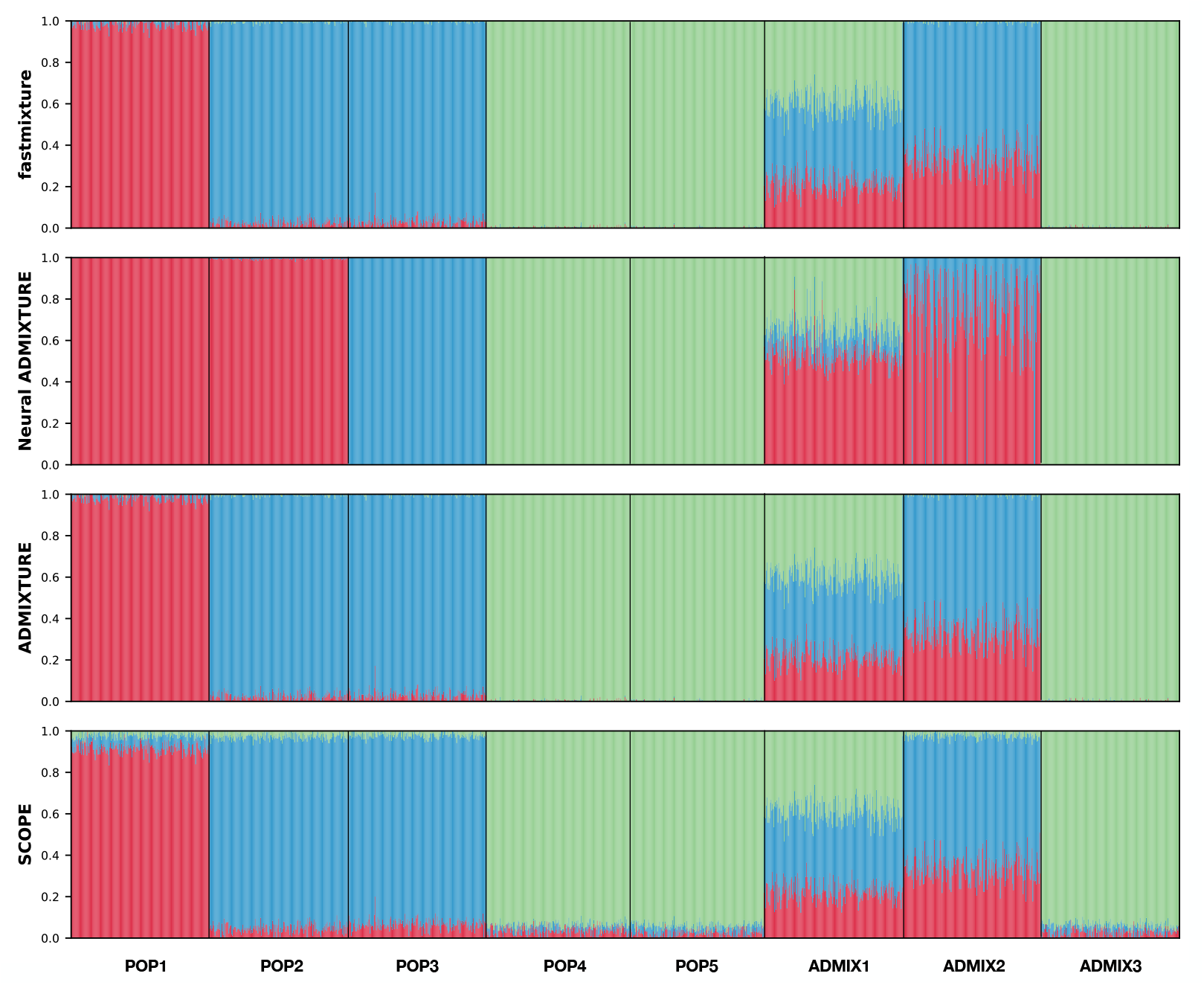
Admixture plots of the ancestry proportions in the simulated individuals for *K* = 3 of *Scenario C* by the different software using their run with the highest log-likelihood.

**Figure S8:**
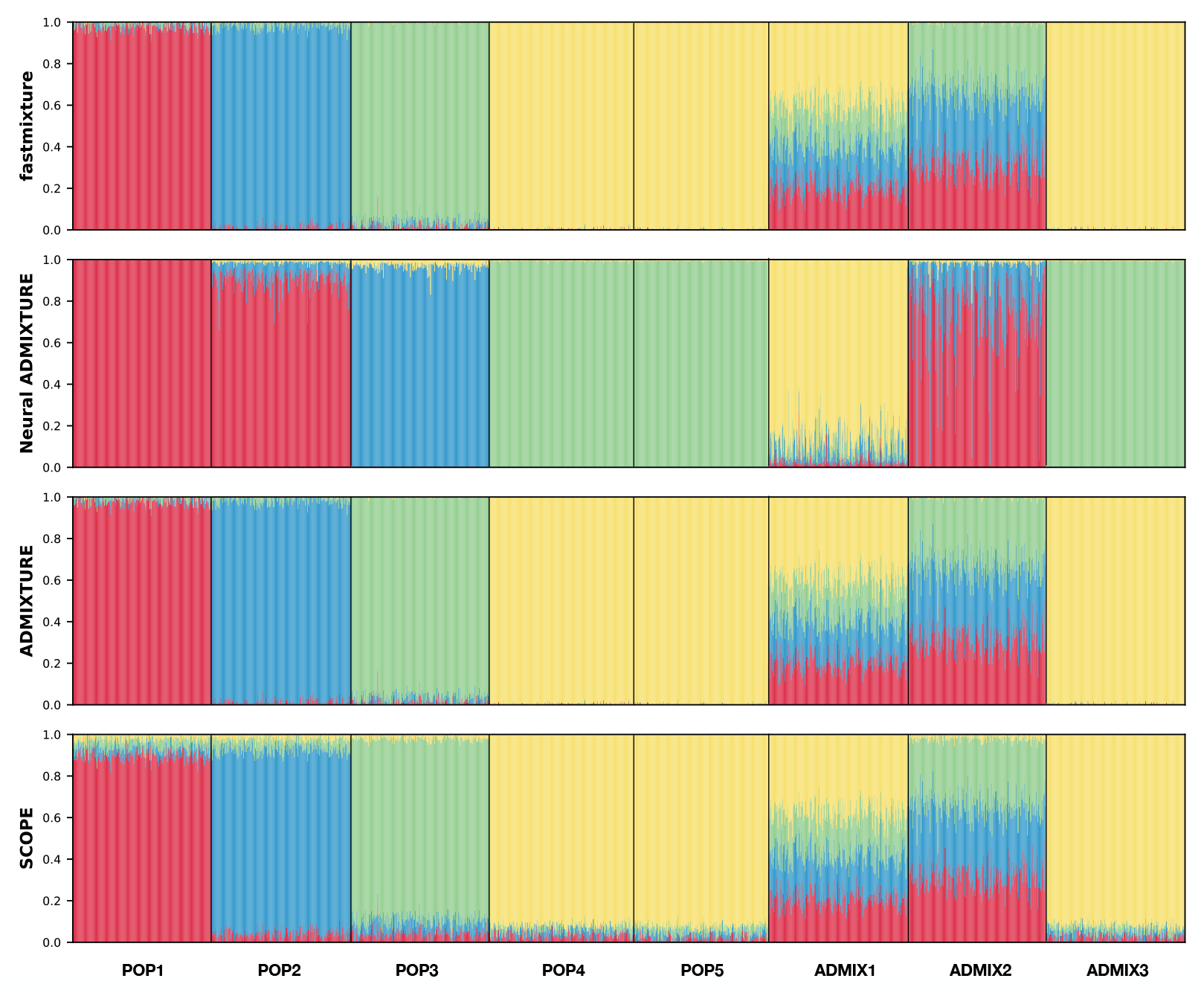
Admixture plots of the ancestry proportions in the simulated individuals for *K* = 4 of *Scenario C* by the different software using their run with the highest log-likelihood.

**Figure S9:**
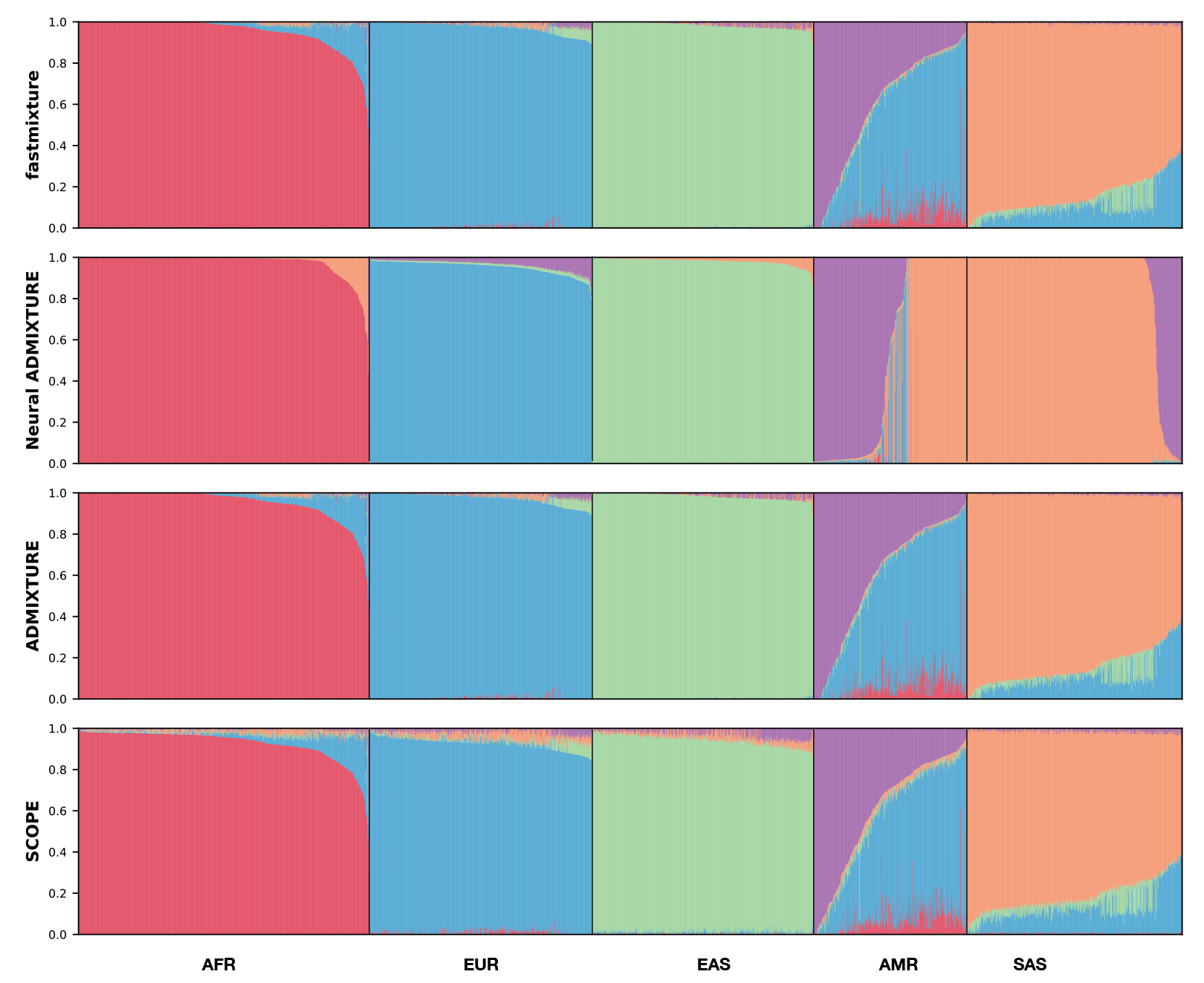
Admixture plots of the estimated ancestry proportions for *K* = 5 in the downsampled version of the 1000 Genomes Project dataset (1KGP Down) by the different software using their run with the highest log-likelihood. AFR: African, EUR: European, EAS: East Asian, AMR: American, and SAS: South Asian ancestry.

**Table S1:** Computational runtimes (in minutes) for the different simulated and empirical datasets for the evaluated methods. The mean across five different runs is reported with the corresponding standard deviation in parenthesis. All analyses have been performed on a cluster with an Intel(R) Xeon(R) Gold 6152 CPU using 64 threads. ^*\**^`ADMIXTURE` has only been evaluated on a single run for 1KGP due to excessive runtime.

| Dataset | fastmixture | Neural ADMIXTURE | ADMIXTURE | SCOPE |
| --- | --- | --- | --- | --- |
| A | 2.70 (0.23) | 3.97 (0.06) | 77.85 (17.97) | 1.69 (0.08) |
| B | 3.09 (0.23) | 3.98 (0.05) | 105.34 (18.33) | 1.96 (0.10) |
| C | 7.11 (0.63) | 6.09 (0.06) | 202.22 (34.27) | 2.31 (0.49) |
| D | 1.32 (0.19) | 2.97 (0.07) | 36.72 (2.37) | 0.24 (0.01) |
| 1KGP Down | 6.62 (0.42) | 10.44 (0.21) | 275.24 (12.77) | 1.97 (0.25) |
| 1KGP | 82.74 (8.13) | 113.54 (3.59) | 2439.92* | 18.69 (2.27) |

**Table S2:** Jensen-Shannon divergence (JSD) measures for the estimated ancestry proportions in the three different simulated datasets for the evaluated methods against the ground truth. The mean across five different runs is reported with the corresponding standard deviation in parenthesis.

| Scenario | fastmixture | Neural ADMIXTURE | ADMIXTURE | SCOPE |
| --- | --- | --- | --- | --- |
| A | 0.0093 (4.2e-5) | 0.0211 (0.0070) | 0.0094 (1.3e-7) | 0.0252 (3.0e-5) |
| B | 0.0079 (4.5e-5) | 0.2075 (0.0044) | 0.0079 (1.5e-7) | 0.0331 (4.2e-5) |
| C | 0.0082 (4.6e-5) | 0.1957 (0.0072) | 0.0083 (1.4e-7) | 0.0398 (2.9e-5) |
| D | 0.0008 (6.7e-7) | 0.0187 (0.0129) | 0.0008 (4.5e-8) | 0.0058 (1.1e-6) |

**Table S3:** Log-likelihood measures in the different simulated and empirical datasets for the evaluated methods. “1KGP” refers to data from the 1000 Genomes Project, where “1KGP (down)” is the downsampled variant. The mean across five different runs is reported with the corresponding standard deviation in parenthesis. ^*\**^`ADMIXTURE` has only been evaluated on a single run for 1KGP due to excessive runtime.

| Dataset | fastmixture | Neural ADMIXTURE | ADMIXTURE | SCOPE |
| --- | --- | --- | --- | --- |
| A | -637729493.6 (0.5) | -640197951.4 (39763.0) | -637729492.4 (0.5) | -637845454.1 (10.0) |
| B | -629360598.6 (0.1) | -633930570.4 (37694.0) | -629360598.5 (1.6) | -629538330.5 (38.1) |
| C | -1006294346.2 (0.6) | -1013880607.9 (52010.8) | -1006294347.6 (4.1) | -1006604915.2 (244.3) |
| D | -429286152.5 (0.2) | -433792567.4 (627402.9) | -429286155.6 (4.1) | -429536651.1 (25.0) |
| 1KGP Down | -1509950457.3 (0.8) | -1531090301.0 (228164.6) | -1509950464.3 (7.5) | -1510431876.5 (254.2) |
| 1KGP | -15097015793.7 (4.9) | -15342582278.2 (3449977.8) | -15097016096.0* | -15101701104.4 (2807.9) |

**Table S4:**
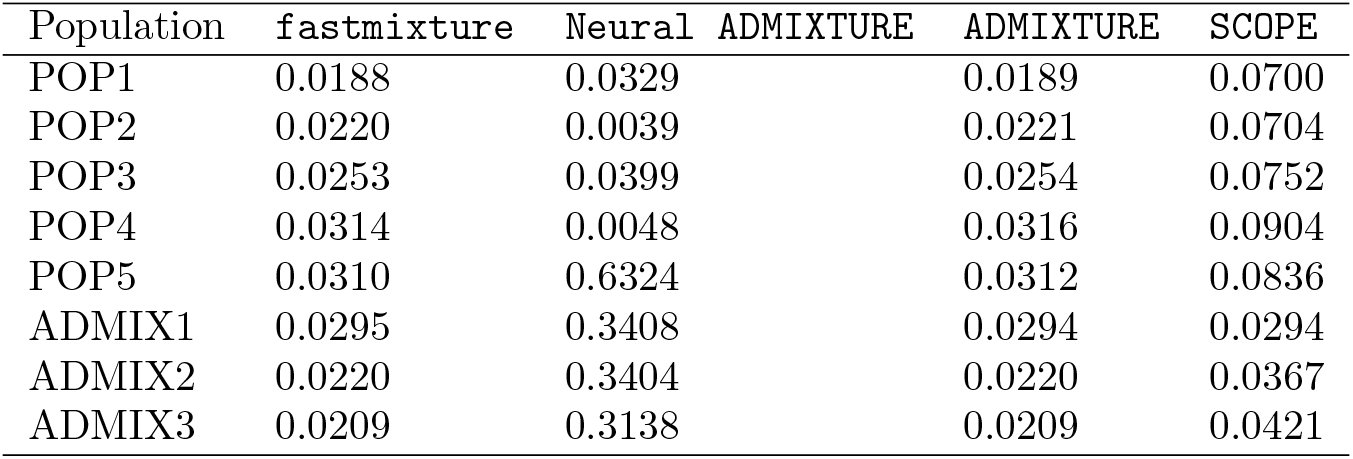
Root mean square error (RMSE) measures for the estimated ancestry proportions in the eight populations of Scenario C for the evaluated methods against the ground truth.

**Table S5:**
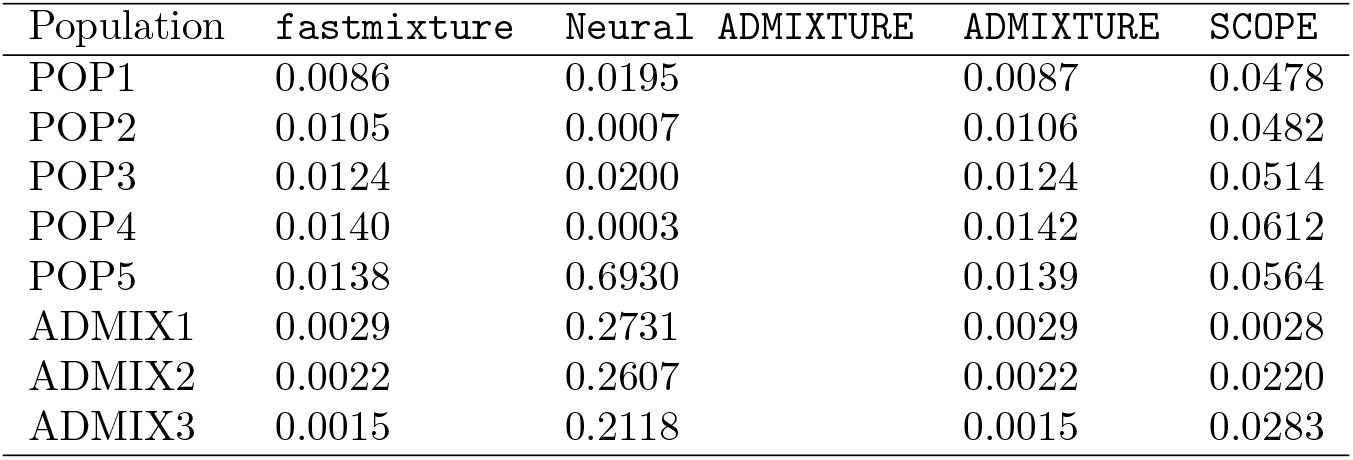
Jensen-Shannon divergence (JSD) measures for the estimated ancestry proportions in the eight populations of Scenario C for the evaluated methods against the ground truth.

**Table S6:** Assessment measures in the simulated dataset of Scenario C using different initial numbers of batches, *B*, in `fastmixture`. The mean across five different runs is reported with the corresponding standard deviation in parenthesis.

| Mini-batches | Runtime (m) | Log-likelihood | RMSE | JSD |
| --- | --- | --- | --- | --- |
| $B = 8$ | 7.86 (1.18) | -1006294348.7 (1.4) | 0.0258 (4.5e-5) | 0.0084 (2.9e-5) |
| $B = 16$ | 7.11 (0.68) | -1006294345.3 (1.0) | 0.0256 (5.1e-5) | 0.0083 (3.0e-5) |
| $B = 32$ | 7.11 (0.63) | -1006294346.2 (0.2) | 0.0255 (8.7e-6) | 0.0082 (4.9e-6) |
| $B = 64$ | 7.53 (0.80) | -1006294353.9 (1.4) | 0.0254 (2.2e-5) | 0.0082 (2.2e-5) |
| $B = 128$ | 9.82 (2.32) | -1006294360.1 (1.5) | 0.0254 (7.9e-6) | 0.0081 (6.7e-6) |

**Table S7:**
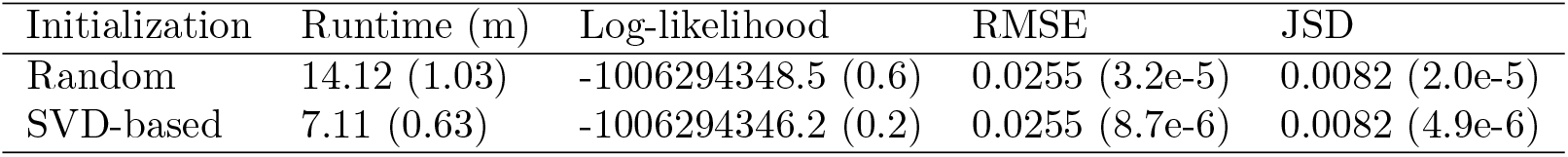
Assessment measures in the simulated dataset of Scenario C using `fastmixture` for two different parameter initialization approaches. The mean across five different runs is reported with the corresponding standard deviation in parenthesis.

| Initialization | Runtime (m) | Log-likelihood | RMSE | JSD |
| --- | --- | --- | --- | --- |
| Random | 14.12 (1.03) | -1006294348.5 (0.6) | 0.0255 (3.2e-5) | 0.0082 (2.0e-5) |
| SVD-based | 7.11 (0.63) | -1006294346.2 (0.2) | 0.0255 (8.7e-6) | 0.0082 (4.9e-6) |

**Table S8:** Log-likelihood measures in the simulated dataset of Scenario C using `fastmixture` for different values of *K*, the number of ancestral populations assumed in the ancestry estimation. The mean across five different runs is reported with the corresponding standard deviation in parenthesis. ^*\**^`ADMIXTURE` had 4/5 runs finding the optimal solution.

|  | **fastmixture** | **Neural ADMIXTURE** | **ADMIXTURE** | **SCOPE** |
| --- | --- | --- | --- | --- |
| $K = 2$ | -1015395537.1 (0.1) | -1016467332.7 (5479.3) | -1015395536.6 (0.0) | -1015464044.1 (0.0) |
| $K = 3$ | -1011420272.3 (0.7) | -1014984382.1 (274833.2) | -1011420270.5 (0.2) | -1011552464.8 (0.1) |
| $K = 4$ | -1008338857.0 (0.4) | -1014255193.2 (127570.6) | -1008546129.6* (463478.4) | -1008524838.1 (4.7) |
| $K = 5$ | -1006294346.2 (0.2) | -1013880607.9 (52010.8) | -1006294347.6 (4.1) | -1006604915.2 (244.3) |

## References

[1] J. Novembre, T. Johnson, K. Bryc, Z. Kutalik, A. R. Boyko, A. Auton, A. Indap, K. S. King, S. Bergmann, M. R. Nelson, et al., “Genes mirror geography within europe,” Nature, vol. 456, no. 7218, pp. 98–101, 2008.

[2] J. Marchini, L. R. Cardon, M. S. Phillips, and P. Donnelly, “The effects of human population structure on large genetic association studies,” Nature genetics, vol. 36, no. 5, pp. 512–517, 2004.

[3] J. K. Pritchard, M. Stephens, and P. Donnelly, “Inference of population structure using multilocus genotype data,” Genetics, vol. 155, no. 2, pp. 945–959, 2000.

[4] H. Tang, J. Peng, P. Wang, and N. J. Risch, “Estimation of individual admixture: analytical and study design considerations,” Genetic Epidemiology: The Official Publication of the International Genetic Epidemiology Society, vol. 28, no. 4, pp. 289–301, 2005.

[5] D. H. Alexander, J. Novembre, and K. Lange, “Fast model-based estimation of ancestry in unrelated individuals,” Genome research, vol. 19, no. 9, pp. 1655–1664, 2009.

[6] B. E. Engelhardt and M. Stephens, “Analysis of population structure: a unifying framework and novel methods based on sparse factor analysis,” PLoS genetics, vol. 6, no. 9, p. e1001117, 2010.

[7] E. Frichot, F. Mathieu, T. Trouillon, G. Bouchard, and O. François, “Fast and efficient estimation of individual ancestry coefficients,” Genetics, vol. 196, no. 4, pp. 973–983, 2014.

[8] J. Meisner and A. Albrechtsen, “Inferring population structure and admixture proportions in low-depth ngs data,” Genetics, vol. 210, no. 2, pp. 719–731, 2018.

[9] I. Cabreros and J. D. Storey, “A likelihood-free estimator of population structure bridging admixture models and principal components analysis,” Genetics, vol. 212, no. 4, pp. 1009–1029, 2019.

[10] A. M. Chiu, E. K. Molloy, Z. Tan, A. Talwalkar, and S. Sankararaman, “Inferring population structure in biobank-scale genomic data,” The American Journal of Human Genetics, vol. 109, no. 4, pp. 727–737, 2022.

[11] P. Gopalan, W. Hao, D. M. Blei, and J. D. Storey, “Scaling probabilistic models of genetic variation to millions of humans,” Nature genetics, vol. 48, no. 12, pp. 1587–1590, 2016.

[12] A. Dominguez Mantes, D. Mas Montserrat, C. D. Bustamante, X. Giŕo-i Nieto, and A. G. Ioannidis, “Neural admixture for rapid genomic clustering,” Nature Computational Science, pp. 1–9, 2023.

[13] A. R. Martin, M. Kanai, Y. Kamatani, Y. Okada, B. M. Neale, and M. J. Daly, “Clinical use of current polygenic risk scores may exacerbate health disparities,” Nature genetics, vol. 51, no. 4, pp. 584–591, 2019.

[14] Y. Wang, K. Tsuo, M. Kanai, B. M. Neale, and A. R. Martin, “Challenges and opportunities for developing more generalizable polygenic risk scores,” Annual review of biomedical data science, vol. 5, no. 1, pp. 293–320, 2022.

[15] H. Zhou, D. Alexander, and K. Lange, “A quasi-newton acceleration for high-dimensional optimization algorithms,” Statistics and computing, vol. 21, pp. 261–273, 2011.

[16] N. Patterson, A. L. Price, and D. Reich, “Population structure and eigenanalysis,” PLoS genetics, vol. 2, no. 12, p. e190, 2006.

[17] S. Ruder, “An overview of gradient descent optimization algorithms,” arXiv preprint arXiv:1609.04747, 2016.

[18] C. C. Chang, C. C. Chow, L. C. Tellier, S. Vattikuti, S. M. Purcell, and J. J. Lee, “Second-generation plink: rising to the challenge of larger and richer datasets,” Gigascience, vol. 4, no. 1, pp. s13742–015, 2015.

[19] C. R. Harris, K. J. Millman, S. J. Van Der Walt, R. Gommers, P. Virtanen, D. Cournapeau, E. Wieser, J. Taylor, S. Berg, N. J. Smith, et al., “Array programming with numpy,” Nature, vol. 585, no. 7825, pp. 357–362, 2020.

[20] Z. Li, J. Meisner, and A. Albrechtsen, “Pcaone: fast and accurate out-of-core pca framework for large scale biobank data,” bioRxiv, pp. 2022–05, 2022.

[21] F. Baumdicker, G. Bisschop, D. Goldstein, G. Gower, A. P. Ragsdale, G. Tsambos, S. Zhu, B. Eldon, E. C. Ellerman, J. G. Galloway, et al., “Efficient ancestry and mutation simulation with msprime 1.0,” Genetics, vol. 220, no. 3, p. iyab229, 2022.

[22] G. Tsambos, J. Kelleher, P. Ralph, S. Leslie, and D. Vukcevic, “link-ancestors: fast simulation of local ancestry with tree sequence software,” Bioinformatics Advances, vol. 3, no. 1, p. vbad163, 2023.

[23] I. H. Consortium et al., “A second generation human haplotype map of over 3.1 million snps,” Nature, vol. 449, no. 7164, p. 851, 2007.

[24] S. Gravel, B. M. Henn, R. N. Gutenkunst, A. R. Indap, G. T. Marth, A. G. Clark, F. Yu, R. A. Gibbs,. G. Project, C. D. Bustamante, et al., “Demographic history and rare allele sharing among human populations,” Proceedings of the National Academy of Sciences, vol. 108, no. 29, pp. 11983–11988, 2011.

[25] S. R. Browning, B. L. Browning, M. L. Daviglus, R. A. Durazo-Arvizu, N. Schneiderman, R. C. Kaplan, and C. C. Laurie, “Ancestry-specific recent effective population size in the americas,” PLoS genetics, vol. 14, no. 5, p. e1007385, 2018.

[26] G. P. Consortium et al., “A global reference for human genetic variation,” Nature, vol. 526, no. 7571, p. 68, 2015.

